# Conditional, inducible gene silencing in dopamine neurons reveals a sex-specific role for Rit2 GTPase in acute cocaine response and striatal function

**DOI:** 10.1101/658856

**Authors:** Carolyn G. Sweeney, Patrick J. Kearney, Rita R. Fagan, Lindsey A. Smith, Nicholas C. Bolden, Rubing Zhao-Shea, Iris V. Rivera, Jenya Kolpakova, Jun Xie, Guangping Gao, Andrew R. Tapper, Gilles E. Martin, Haley E. Melikian

## Abstract

Dopamine (DA) signaling is critical for movement, motivation, and addictive behavior. The neuronal GTPase, Rit2, is enriched in DA neurons (DANs), binds directly to the DA transporter (DAT), and is implicated in several DA-related neuropsychiatric disorders. However, it remains unknown whether Rit2 plays a role in either DAergic signaling and/or DA-dependent behaviors. Here, we leveraged the TET-OFF system to conditionally silence Rit2 in *Pitx3^IRES2-tTA^* mouse DANs. Following DAergic Rit2 knockdown (Rit2-KD), mice displayed an anxiolytic phenotype, with no change in baseline locomotion. Further, males exhibited increased acute cocaine sensitivity, whereas DAergic Rit2-KD suppressed acute cocaine sensitivity in females. DAergic Rit2-KD did not affect presynaptic TH and DAT protein levels in females, nor was TH was affected in males; however, DAT was significantly diminished in males. Paradoxically, despite decreased DAT levels in males, striatal DA uptake was enhanced, but was not due to enhanced DAT surface expression in either dorsal or ventral striatum. Finally, patch recordings in nucleus accumbens (NAcc) medium spiny neurons (MSNs) revealed reciprocal changes in spontaneous EPSP (sEPSP) frequency in male and female D1+ and D2+ MSNs following DAergic Rit2-KD. In males, sEPSP frequency was decreased in D1+, but not D2+, MSNs, whereas in females sEPSP frequency decreased in D2+, but not D1+, MSNs. Moreover, DAergic Rit2-KD abolished the ability of cocaine to reduce sEPSP frequency in D1+, but not D2+, male MSNs. Taken together, our studies are among the first to acheive AAV-mediated, conditional and inducible DAergic knockdown *in vivo*. Importantly, our results provide the first evidence that DAergic Rit2 expression differentially impacts striatal function and DA-dependent behaviors in males and females.

## Introduction

Dopamine (DA) neurotransmission is critical for learning, motivation, movement, anxiety, and rewarding behaviors [1,2]. DAergic dysfunction has profound clinical consequences, as evidenced by multiple DA-related neurological and neuropsychiatric disorders, including Parkinson’s disease (PD), attention-deficit hyperactivity disorder (ADHD), schizophrenia, autism spectrum disorder (ASD), and anxiety [3–6]. DAN cell bodies are densely clustered within the midbrain structures of the substantia nigra pars compacta (SNc) and ventral tegmental area (VTA). The SNc primarily projects to the dorsal striatum (DS), whereas the VTA densely innervates the prefronal cortex (PFC) and NAcc. DA is largely modulatory in these regions, where it significantly modulates target cell excitability. DA release in the NAcc is required for the rewarding behaviors elicited by addictive drugs. Moreover, addictive psychostimulants, such as cocaine and amphetamines, increase NAcc DA levels by inhibiting the presynaptic DA clearance by the plasma membrane [7]. Thus, mechanisms that alter DAergic signaling are predicted to have a broad downstream impact on both neuronal excitability and response to addictive drugs.

The neuronal, Ras-like GTPase, Rit2 (aka: Rin, **<u>r</u>**as-like **<u>i</u>**n **<u>n</u>**eurons), is enriched in DANs [8] and is required for NGF-mediated MAP kinase activation in neuroendocrine cells [9,10]. Multiple genome-wide association studies (GWAS) identified Rit2 in several DA-related disorders, including PD [11], ASD [12], essential tremor, schizophrenia, and bipolar disorder [13–15]. We previously demonstrated that Rit2 binds directly to the presynaptic, cocaine-sensitive DAT, and that Rit2 activity is required for PKC-stimulated DAT endocytosis in cell lines [16]. Despite the putative linkage between Rit2 and DAergic function, it is entirely unknown whether or how Rit2 influences either DAergic function and/or DA-related behaviors.

In the current study, we aimed to directly test a putative role for Rit2 in DA-dependent behaviors and striatal physiology, by conditionally silencing Rit2 in DANs. Conditional, inducible, rAAV-mediated gene delivery is an indispensable tool to define neuronal circuitry and pinpoint molecular mechanisms required for neuronal development, signaling and dysfunction. However, *in vivo* conditional gene expression approaches, such as Cre-lox system or cell-specific promotors, are not ideal for DAergic, AAV-mediated shRNA expression. First, the hairpin-like secondary structures, inherent to shRNAs and lox sites, create AAV genome instability that heterogeneously truncates viral genomes [17]. Moreover, self-complimentary AAVs have packaging size constraints that prohibit utilizing large DAergic promoters, such as those for tyrosine hydroxylase (TH) and DAT, to conditionally drive shRNA expression [18]. Here, we leveraged the TET-OFF system to acheive AAV-mediated, conditional and inducible gene silencing in adult mouse DANs, and utilized this approach to test whether Rit2 expression is required for DA-dependent behaviors and striatal function.

## Materials and Methods

### cDNA constructs

Mouse Rit2 (mRit2) cloned into pGEX2T was a gift from Julian Downward (Addgene plasmid #55663)[59]. To generate RFP-mRit2, the mRit2 coding region was PCR-amplified and subcloned in-frame into the pTagRFP-C vector (Evrogen) at HindIII/XbaI sites. Rit2 shRNAs and pGIPZ controls were obtained from Dharmacon. Mature antisense sequences for each shRNA are as follows: shRit2#1 (clone V3LMM_441839) TTATCTTCTTCCACAGGCT and shRit2#2 (clone V3LMM_441840) TCATAGGTGTGACGGACCT.

### Antibodies

Primary antibodies used: Rat anti-DAT (MAB369), rabbit anti-TH (AB152; IHC) and mouse anti-TH (clone LCN1; immunoblots) were from Millipore Sigma. Chicken anti-GFP (PA1-9533) and mouse anti-RFP (clone RF5R) were from Thermo-Fisher, and mouse anti-actin (clone SPM161) was from Santa Cruz. HRP-conjugated secondary antibodies: Goat anti-rat was from Millipore and Jackson Immuno Research, and goat anti-mouse was from Jackson ImmunoResearch). AlexaFluor-conjugated, cross-absorbed secondary antibodies were from Jackson ImmunoResearch.

### AAV constructs and viral production

*pscAAV-TRE-eGFP and pscAAV-TRE-miR33-shRNA-eGFP plasmids*. pTre3G promoter was isolated from pTre3G (Clone tech) plasmid using EcoRI and SalI. The CB6 promoter in the pscAAV-CB6-eGFP was removed by MluI and BstXI digestion and replaced with the pTre3G promoter to generate pscAAV-TRE3g-eGFP plasmid. pscAAV-TRE3g-eGFP plasmid was digested with BglII and PstI as backbone, miR-33-shRNA was synthesized as insert to generate pscAAV-TRE3g-miR33-shRit2-eGFP. AAV particles (AAV9 serotype) were produced, purified, and titers determined by the University of Massachusetts Medical School Viral Vector Core, as previously described [60].

### Cell culture and transfections

5×10^5^ HEK293T cells/well were seeded in 6 well plates 16-24 hours prior to transfection. Cells were transfected with Lipofectamine 2000 (Thermo Fisher) per manufacturers protocols, using 2μg plasmid DNA/well, at a 2:1 lipid to cDNA ratio. For initial Rit2 shRNA screens: cells were co-transfected with RFP-Rit2 reporter (0.67μg/well) and either shRit2#1, shRit2#2, or pGIPZ vector control (1.33μg/well). For pAAV validation: 660ng shRNA plasmid, 330ng RFP-mRit2 reporter plasmid, and 1 μg rtTA plasmid, and cells incubated ±500ng/mL doxycycline, 48 hrs, 37°C.

### Mice

*Pitx3^IRES-tTA^*/+ were the generous gift of Dr. Huaibin Cai (National Institute on Aging), and were continually backcrossed to *C57Bl/6* mice. Homozygous *Pitx3^IRES-tTA^* were crossed to *D1-tD-tomato* mice (Dr. Gilles Martin) to generate *Pitx3^IRES-tTA^*; *D1-tD-tomato* mice used for electrophysiology studies. Mice were maintained in 12hr light/dark cycle at constant temperature and humidity. Food and water was available ad libitum, and mice were maintained on either standard or doxycycline-supplemented (200mg/kg) chows (S3888, BioServ), as indicated. All studies were conducted in accordance with UMASS Medical School IACUC Protocol A-1506 (H.E.M).

### Stereotaxic viral delivery

Adult mice (minimum 3 weeks age) were anesthetized with 100mg/kg ketamine (Vedco Inc.) and 10mg/kg xylazine (Akorn Inc.), I.P. 20% mannitol (NeogenVet) was administered i.p. 15 minutes (minimum) prior to viral delivery, to increase viral spread [61]. Mice heads were shaved and placed in the stereotaxic frame (Stoelting Inc.). 1μl of the indicated viruses were administered bilaterally into the VTA (Bregma: anterior/posterior: −3.08mm, medial/lateral: ±0.5mm, dorsal/ventral: −4.7mm) at a rate of 0.2μL/min over 5 minutes. Syringes were left in place for a minimum of 3 minutes post-infusion prior to removal. Mice were individually housed, for a minimum of four weeks (biochemistry and electrophysiology) or six weeks (behavior) before experiments were performed. Dox-treated mice were placed on dox-supplemented chow, at minimum, 4 days prior to viral injection, and were maintained on (+)dox chow throughout recovery and experimental procedures. For (−)dox mice, viral injections were confirmed by posthoc immunohistochemistry, validating midbrain GFP expression.

### Midbrain tissue isolation and RT-qPCR

Mice were sacrificed, brains were rapidly removed and either flash frozen on dry ice and sectioned on a cryostat or were sectioned fresh on a Vibratome in ice cold cutting solution (2.5mM KCl, 1.25mM NaH_2_PO_4_, 20mM HEPES, 2mM thiourea, 5mM sodium ascorbate, 3mM sodium pyruvate, 92mM N-methyl-D-glucamine, 30mM NaHCO_3_, 25mM D-glucose). For frozen tissue: 10μm coronal sections were made using cryostat (Leica Microsystems Inc.). A Veritas Microdissection System Model 704 (Arcturus Bioscience) was used to microdissect SNc/VTA- or SNr-enriched regions. For fresh tissue: 200μm sections were made, and 1mm diameter tissue punches were made in VTA/SNc or SNr, under magnification. Total RNA was isolated using RNAqueous kit (Thermo) per manufacturers protocol. Following DNAase treatment (30 min, 37°C), cDNA was generated (Retroscript; Thermo) and mRNA levels were quantified by RT-qPCR, using an Applied Biosystems 7500 Real-Time System and TaqMan assays: Rit2 (Mm01702749_mH, GAPDH: Mm99999915_g1, DAT: Mm00438388_m1, TH: Mm00447557_m1 (Applied Biosystems). Each independent mRNA sample was analyzed in triplicate, and expression levels were normalized to GAPDH expression, using 2^−ΔCt^ method [62].

### Immunohistochemistry

Mice were transcardially perfused with ice-cold phosphate-buffered saline, pH 7.4 (PBS), followed by ice cold 4% paraformaldehyde/PBS (w/v). Brains were removed and submerged in 4% PFA for 24hr 4°C, then dehydrated in 30% sucrose/PBS (w/v), 3-7 days 4°C. Brains were flash frozen on dry ice and 25μm coronal sections were prepared on a microtome. Tissue was blocked and permeabilized in IHC blocking solution (5% normal goat serum, 1% H_2_O_2_, 0.1% Triton-X 100 in PBS), 1 hour, RT. All antibodies were diluted in IHC blocking solution at the following dilutions: rabbit anti-TH (1:500), chicken anti-GFP (1:500), goat anti-rabbit AlexaFluor594 (1:250), goat anti-chicken AlexaFluor488 (1:250). Slices were co-incubated in the indicated primary antibodies, 2 days, 4°C, washed and co-incubated with the indicated secondary antibodies, 2 hours, RT. Slices were washed, mounted on glass slides, and air-dried prior to mounting on glass coverslips in Prolong Gold Anti-fade with DAPI (Thermo-Fisher). Slides cured overnight and samples were visualized with a Zeiss Axiovert 200M microscope, using a 5X, N.A.0.15 Plan-Neofluar objective. Images were captured using a Retiga-R1 CCD camera (QImaging) and Slidebook 6 software (Intelligent Imaging Innovations), using identical intrachannel exposure times across all samples. Images were processed using Photoshop, and identical red and green levels were applied across samples, within each imaged brain region.

### Striatal Slice preparation

*Ex vivo* striatal slices were prepared 3-4 weeks following viral injection. For electrophysiology slices, mice (P49-P56) were anesthetized with isoflurane and were transcardially perfused with ice-cold NMDG cutting solution (in mM: 20 HEPES, 2.5 KCl, 1.25, NaH_2_PO_4_, 30.0 NaHCO_3_, 25.0 glucose, 0.5 CaCl_2_·4H_2_O, 10.0 MgSO_4_·7H_2_O, 92 N-methyl-D-glucamine (NMDG), 2.0 thiourea, 5.0 Na^+^-ascorbate, 3.0 Na^+^-pyruvate) followed by rapid decapitation. For biochemical and uptake assays, mice were sacrificed by cervical dislocation followed by rapid decapitation. Brains were removed and 300μm coronal slices were prepared with a VT1200 Vibroslicer (Leica) in ice-cold, oxygenated NMDG cutting solution. Hemislices were prepared by bisecting slices at the midline, and were recovered as follows: For biochemical and uptake studies, slices recovered in oxygenated aCSF (in mM: 125 NaCl, 2.5 KCl, 1.24, NaH_2_PO_4_, 26 NaHCO_3_, 11.0 glucose, 2.4 CaCl_2_·4H_2_O, and 1.2 MgCl_2_·6H_2_O), 40 min, 31°C; for patch recordmings, slices recovered in NMDG cutting solution, 20 min, 31°C, followed by 1 hour (minimum) at room temperature in oxygenated aCSF.

### Immunoblots

Slices and cultured cells were solubilized in RIPA (10mM Tris base pH 7.4, 150mM NaCl, 1mM EDTA, 0.1% (v/v) SDS, 1% (v/v) Triton-X 100, 1% (w/v) sodium deoxycholate) supplemented with protease inhibitors (1mM PMSF, 1μg/mL leupeptin, 1μg/mL pepstatin, 1μg/mL aprotinin) for a minimum of 30 minutes at 4°C. For brain slices, lysates were additionally triturated sequentially through a 200μL pipet tip, followed by 22G and 26G tech-tips [63], and insoluble material was pelleted by centrifugation (14,000 x g, 20 min, 4°C). Protein concentrations were determined by BCA protein assay (Thermo Fisher) using BSA as a standard. The indicated amounts of protein were incubated with 2x SDS-PAGE sample buffer and rotated, 30 minutes, RT. Samples were resolved by SDS-PAGE and were assessed by immunoblot using the indicated antibodies at the following dilutions: anti-DAT (1:2000), anti-actin (1:10,000), anti-TH (1:5000), anti-RFP (1:2000). Immunoreactive bands were captured using a VersaDoc Digital Imaging station, and non-saturating signals were quantified using Quantity One software (BioRad).

### Ex vivo slice [^3^H]DA uptake

Following their preparation and recovery, hemislices were transferred to 12-well plates fitted with cell culture inserts and containing oxygenated aCSF, 37°C. All slices were pretreated with 100nM desipramine, 30 min, 37°C to block potential uptake contribution from the norepinephrine transporter, and hemislices used to define non-specific DA accumulation were additionally pretreated with 10μM GBR12909. DA transport was initiated by adding 1μM [^3^H]DA in aCSF supplemented with 10μM each pargyline and sodium L-ascorbate, and proceeded for 10min, 37°C. Uptake was terminated by rapidly washing slices with ice-cold aCSF, followed by three 5 min incubations in ice-cold aCSF. Slices were lysed in RIPA buffer containing protease inhibitors, 30 min, 4°C, were triturated sequentially through a 200μL pipet tip, followed by 22G and 26G tech-tips [63], and insoluble material was removed by centrifugation (18,000 x g, 10 min, 4°C). Protein concentrations were determined using the BCA protein assay (Pierce) and [^3^H]DA signals were quantified by liquid scintillation counting from three technical replicates per hemi-slice (100 μg total lysate per replicate). Specific DAT-mediated uptake was determined by subtracting the accumulated counts from contralateral, GBR12909-treated hemislices. To control for normal rostral-caudal variations in DAergic terminal density, TH content in each hemi-slice was determined by immunoblotting a 20μg lysate aliquot, and was normalized to actin loading controls. DA uptake in each hemi-slice was subsequently normalized to the TH/actin value.

### Ex vivo slice biotinylation

Following initial recovery, hemi-slices were were transferred to 12-well plates fitted with cell culture inserts, incubated for 30 min, 37°C in oxygenated aCSF, and surface DAT was quantified by surface biotinylation as previously described by our laboratory [19,20]. Specifically, slices were transferred to ice baths and surface proteins were labeled with membrane-impermeant sulfo-NHS-SS-biotin (1.0mg/ml), 45min, 0°C. Residual reactive biotin groups were quenched with ice-cold aCSF, supplemented with 100mM glycine (2 x 20 min incubations), followed by three rapid washes with ice-cold aCSF. Hemi-slices were enriched for dorsal and ventral striatum, by sub-dissecting in a line from the anterior commissure to the lateral olfactory tract. Sub-dissected slices were lysed in RIPA buffer containing protease inhibitors, and tissue was disrupted by triturating sequentially through a 200μL pipet tip, followed by 22G and 26G tech-tips [63]. Samples rotated 30 min, 4°C, insoluble material was removed by centrifugation, and protein concentrations were determined using the BCA protein assay (Thermo Fisher). Biotinylated proteins were isolated by batch streptavidin chromatography, overnight with rotation, 4°C, at a ratio of 20μg striatal lysate to 30μL streptavidin agarose beads, which was empirically determined to recover all biotinylated DAT. Recovered proteins were washed with RIPA buffer and eluted from beads in 2X SDS-PAGE sample buffer, 30min, room temperature with rotation. Eluted (surface) proteins and their respective lysate inputs were resolved by SDS-PAGE, and DAT was detected by immunoblotting as described above. %DAT at the cell surface was determined by normalizing surface signals to the corresponding total DAT input signal in a given hemi-slice, detected in parallel.

### Mouse Behavior

For all tests, mice were habituated to the experimental room for >30 minutes. Rooms were kept in dim light, and white noise was used to maintain constant ambient sound.

### Open field test

Mouse activity was individually assessed in a 38.5×40cm open arena, 10 minutes Total distance traveled, and time spent in the arena center vs. periphery, were measured with EthoVision software (Noldux).

### Elevated Plus Maze (EPM)

Mice were placed in the maze junction and their position over five minutes was recorded by MED-PC IV software (MED Associates, Inc.). The time spent in open and closed arms, in the junction, and total entries were determined.

### Cocaine induced locomotor activity

Ambulatory activity was recorded in individual photobeam activity chambers (San Diego Instruments), following the indicated drug injections. On all test days, mice habituated to the chamber for 45 minutes and postinjection locomotor activity was recorded for 90 minutes. On day one, mice received saline (10mL/kg, I.P.). On day two, mice received a single cocaine injection (5, 15, or 30 mg/kg, I.P.). For data analysis, total horizontal movement was measured, and mouse movement in 5-minute bins on cocaine-treatment days was normalized to the corresponding 5-minute bin from saline-treatment days for each mouse.

### Electrophysiology

Slices were transferred to a submersion recording chamber and were perfused with a oxygenated aCSF, at a constant rate of 1–2 mL/min. NAcc MSNs were visualized in infrared differential interference contrast videomicroscopy using a fully motorized microscope mounted with 10x and 60x objective (Olympus Microscopy), and td-Tomato+, D1+ MSNs were identified by fluorescence microscopy. We acquired current and voltage traces using borosilicate glass electrodes (1.5 mm OD, 4–6 MΩ resistance) filled with an internal solution containing (in mM): 120 K-methanesulfonate, 20 KCl, 10 HEPES, 2 K_2_ATP, 2 K_2_GTP, and 12 phosphocreatine. We acquired and filtered EPSPs at 10 kHz and 1.2 kHz, respectively, with an EPC-10 amplifier and Patchmaster acquisition software (HEKA). To measure MSNs membrane input resistance we injected a 800 ms-long negative current step (−100 pA). To assess action potential (AP) threshold, AP overshoot, and fast after hyperpolarization (AHP), we evoked action potentials with the following ramp protocol:, a current step of −120 pA was initially injected for 10 ms before reaching a maximum value of +250 pA in 800 ms. This protocol was repeated 4 consecutive times and the properties of the first action potential of each trace were measured. We recorded 4 consecutive 1 min long gap-free spontaneous excitatory postsynaptic potentials (sEPSPs) in presence of 15μM bicuculline, followed by bath perfusion with 10□M cocaine, at resting membrane potential (−85 ± 1.2 mV). The final minute of each condition was used for analysis. Recordings showing a rundown of sEPSPs frequency were rejected. We analyzed sEPSPs amplitude and frequency with Clampfit (pClamp 11 software suite, Molecular Devices, CA). We monitored series resistance by comparing EPSPs decay time before and after induction using Clampfit event template analysis. We rejected recordings with changes of resting potential larger than 1 mV.

### Statistical Analysis

All data were analyzed using GraphPad Prism software. Statistical outliers within data sets were determined prior to analysis, using Rout’s test. Specific statistical tests used are detailed within each figure legend. Comparisons between two experimental conditions were made using either Student’s t test, or KS test for cumulative probabilities. Significance across multiple experimental samples was determined by either one-way or two-way ANOVA, as indicated, with posthoc analyses to determine significance between stated experimental groups.

## Results

### Conditional and inducible DAergic Rit2 knockdown using the Tet-OFF/ON system

We identified two mouse-specific, Rit2-directed candidate shRNAs, which significantly reduced RFP-mRit2 protein *in vitro* (Fig. S1a). We chose the more efficacious shRNA (#1) for *in vivo* studies, and cloned it into an artificial microRNA backbone within the pscAAV-TRE-mIR33-eGFP vector (hereafter referred to as shRit2). When coexpressed with rtTA (TET-ON), shRit2 efficaciously silenced RFP-mRit2 in a doxycycline-dependent manner (Fig. S1b). We therefore used this plasmid to generate AAV9 particles for *in vivo* Rit2 knockdown (Rit2-KD) studies.

### AAV-mediated Rit2 knockdown in Pitx3^IRES-tTA^/+ mouse dopaminergic neurons

We used *Pitx3^IRES2-tTA^* mice, which selectively expresses tTA in TH+ midbrain neurons [21], to conditionally drive DAergic TRE-shRit2 expression (see schematic, Fig. 1a). To test whether shRit2 could achieve conditional, inducible, and efficacious *in vivo* Rit2-KD, we bilaterally injected *Pitx3^IRES2-tTA^*/+ VTA with AAV9-TRE-shRit2 (Fig. 1b), and maintained mice ±200mg/kg doxycycline diet for 6 weeks post-injection. In (−)dox mice, GFP reporter expression was clearly visible in TH+ cell bodies within the VTA and SNc (Fig. 1c, e), but not in adjacent TH^−^ cells of the interpeduncular nucleus (IPN; Fig. 1c) and substantia nigra reticulata (SNr; Fig. 1e). In contrast, in (+)dox mice, GFP expression was markedly suppressed in TH+ neurons (Fig. 1d,f), consistent with dox-mediated suppression of TRE-driven shRNA and GFP expression. RT-qPCR studies revealed that AAV9-TRE-shRit2 did not affect Rit2 expression in the non-DAergic SNr (Fig. 1g), but significantly decreased Rit2 expression in SNc-enriched tissue from females (Fig. 1h), and males (Fig. 1i), as compared to control (+dox) mice. shRit2 was highly specific for Rit2, as expression of Rit1, the closest ras homolog to Rit2, was unaffected by shRit2 (Fig. S1c,d).

**Figure 1.**
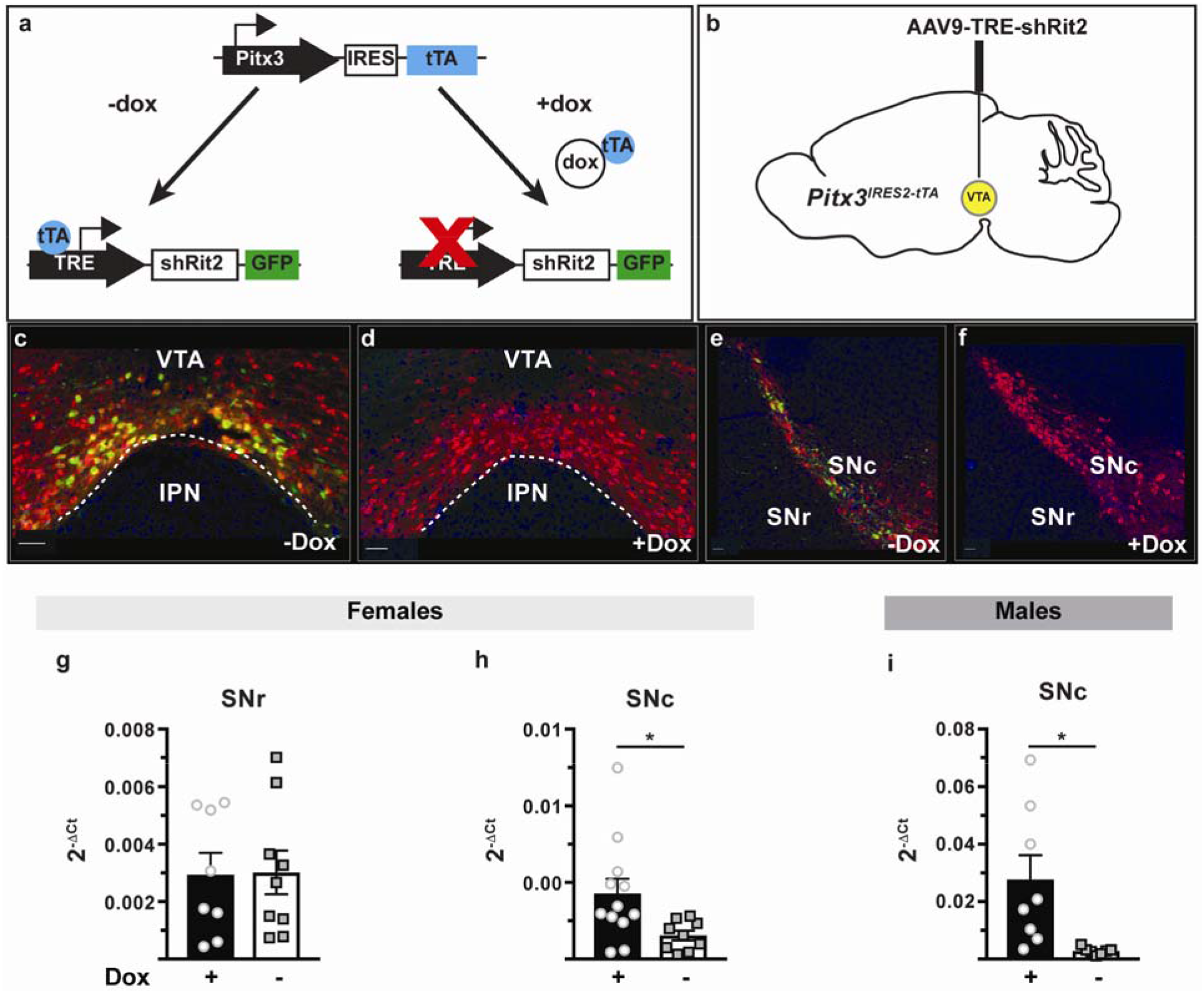
Conditional and inducible gene silencing in midbrain DA neurons. **(a)** *Strategy for conditional and inducible Rit2 silencing in DA neurons*. Conditional and inducible Rit2-KD in DA neurons was achieved using the Tet-Off system, in which tTA expression was driven downstream of the DAergic transcription factor, Pitx3. Following injection with an AAV encoding TRE-shRit2-GFP, tTA binds the Tet-response element (TRE) and drives both shRit2 and GFP reporter expression under (−)dox conditions. When mice are maintained on a (+)dox diet, doxycycline binds the tTA and prevents TRE-driven shRit2 and GFP expression. **(b)** *Pitx3^IRES-tTA^*/+ mouse VTA was bilaterally injected with AAV9-TRE-GFP and mice were maintained ±dox diet ≥6 weeks. **(c-f)** *Immunohistochemistry:* Brains were sectioned 6 weeks postinjection, stained for TH (red) and GFP (green), and imaged and imaged as described in *Methods*. GFP was selectively expressed in TH+ midbrain neurons, and (+)dox diet suppressed GFP expression in both the VTA **(c,d)** and SNc **(e,f)**. **(g-i)** *RT-qPCR studies*. The indicated brain regions were harvested by tissue punch and/or laser capture microscopy, and Rit2 mRNA levels were quantified by RT-qPCR, as described in *Methods*. Data are presented as 2^−ΔCt^ values, ±S.E.M **(g, h)** Female data: AAV9-TRE-shRit2 did not affect Rit2 expression in non-DAergic, SNr (**g**; p=0.92), but significantly decreased Rit2 expression in SNc (**h**; *p<0.02). **(i)** Male data from midbrain punches: AAV9-TRE-shRit2 significantly decreased midbrain Rit2 (p=0.021), one-tailed Student’s t test, n=7-12.

### DAergic Rit2 knockdown impacts baseline anxiety, but not locomotion

DAergic transmission is central to multiple rodent behaviors, including locomotion [22] and anxiety [23]. We first assessed mice in the in the open field test (OFT) to test whether DAergic Rit2-KD affected baseline locomotion. We observed no significant effect of sex on baseline locomotion (p=0.94, two-way ANOVA, sex effect, F_1,26_=0.058, n=7-8), therefore we grouped sexes for OFT analyses. DAergic Rit2-KD (−dox) did not affect total distance traveled, as compared to controls (+dox) (Fig. 2a). However, Rit2-KD mice spent significantly more time in the arena center than controls (Fig. 2b), suggesting an anxiolytic phenotype. We further tested this possibility using the elevated plus maze (EPM). Similar to OFT results, sex did not affect time spent in open arms (p=0.63, two-way ANOVA, sex effect, F_1,26_=0.24, n=7-8); therefore, we grouped both sexes for analysis. Rit2-KD (−dox) mice spent significantly more time in the open arms as compared to controls (+dox; Fig. 2c), consistent with an anxiolytic effect. There was no significant difference in either the time spent in the closed arms (p=0.35; Fig. S2a) or in the junction (p=0.13; Fig. S2b); however Rit2-KD significantly increased total entries (Fig. S2c). The observed anxiolytic effect was not due to the dox diet (Fig. S2d). Taken together, these data demonstrate that DAergic Rit2 expression impacts baseline anxiety-like behaviors in both sexes, but does not impact baseline locomotion.

**Figure 2.**
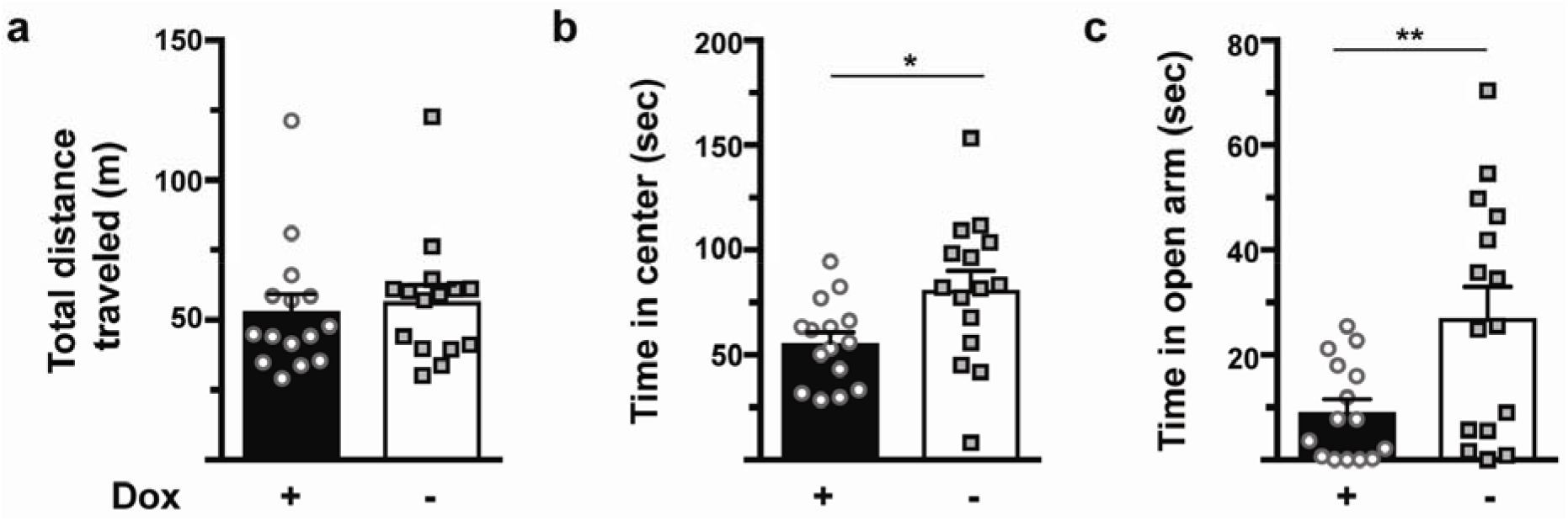
Rit2-KD in DA neurons does not affect baseline locomotor activity, but reduces generalized anxiety. AAV9-TRE-shRit2 was injected into male and female *Pitx3^IRES2-tTA^*/+ VTA, and mice were maintained ±dox diet for 6 weeks prior to assessing behavior. Male and female data were pooled, and −dox vs. +dox results were compared using a two-tailed Student’s t test (n=15). **(a, b)** *Open field test*. **(a)** Total distance traveled (p=0.67). **(b)** Time spent in center of field. *p<0.02 **(c)** *Elevated plus maze*. Rit2-KD (−dox) mice spent significantly more time in open arms than control (+dox) mice. *p<0.01.

### DAergic Rit2 knockdown differentially impacts acute locomotor response to cocaine in males and females

Cocaine is a potent DAT inhibitor that acutely increases extracellular DA levels and elicits robust hyperlocomotion [7]. Therefore, we used shRit2 to test whether DAergic Rit2 expression is required for cocaine-induced hyperlocomotion following a single, I.P. cocaine injection (5, 15 or 30 mg/kg). In males, DAergic Rit2-KD significantly increased acute cocaine responses over time at all doses tested (−dox; Fig. 3a-c). Male total ambulation was also significantly higher than controls at 5 and 15 mg/kg cocaine (Fig 3a,b *insets*). In contrast, in females, DAergic Rit2-KD significantly decreased acute cocaine responses over time at the 15 mg/kg dose, but not at either the 5 or 30 mg/kg doses (Fig. 3d-f), and female total ambulation following cocaine injection was also significantly lower than controls at the 15 mg/kg dose (Fig. 3e, *inset*). The observed effects were not due to either the dox treatment or viral injection in either sex at 15mg/kg, the dose where Rit2-KD drove sex-dependent differences in cocaine response (Fig. S3a,b). Moreover, there was no significant effect in female fine motor behavior at any cocaine dose (Fig. S3c), suggesting that the attenuated cocaine response was not likely due to increased stereotypy.

**Figure 3.**
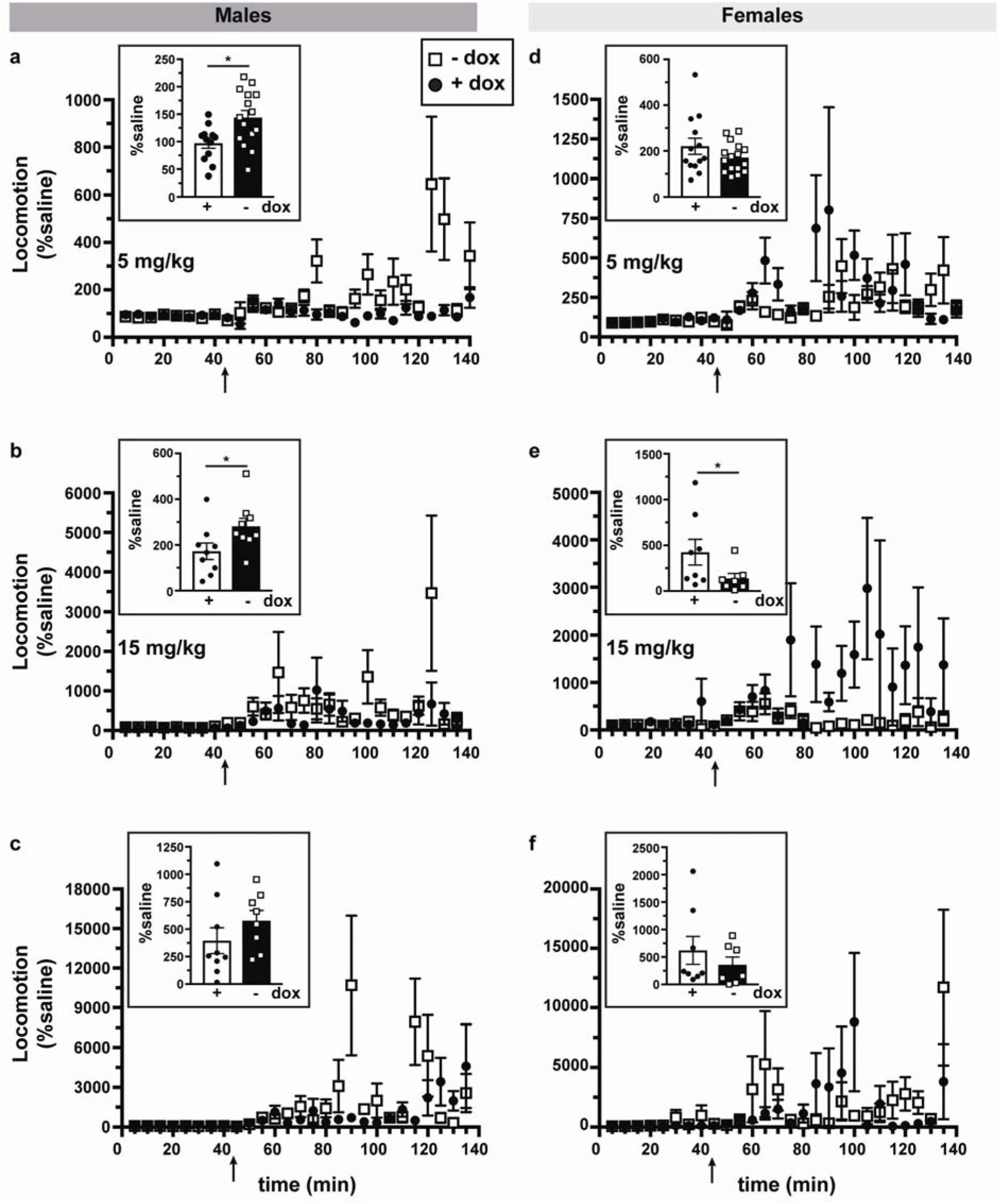
Rit2-KD in DA neurons increases acute cocaine sensitivity in males, and decreases acute cocaine sensitivity in females. *Cocaine-induced hyperlocomotion*. AAV9-TRE-shRit2 was bilaterally injected into male and female *Pitx3^IRES2-tTA^*/+ VTA, and mice were maintained ±dox diet, 6 weeks prior to assay. Horizontal ambulation was recorded in PAS chambers as described in *Materials and Methods*. Average ambulation ±S.E.M is displayed over time, following either a 5, 15, or 30 mg/kg I.P. cocaine injection. Values in each time bin were normalized to the response to a saline injection for each animal, acquired one day prior to cocaine administration. Arrows indicate time of cocaine injection. Timecourse data were analyzed by two-way ANOVA. *Insets:* Total ambulation across the indicated session is presented, and was analyzed using an unpaired, two-tailed Student’s t test. **(a-c)** *Male locomotor activity*. Rit2 knockdown in DANs significantly increased male response to cocaine at all doses tested (5 mg/kg: time: F_(27, 645)_ = 3.475, p<0.001, knockdown: F_(1, 645)_ = 26.1, p<0.001; 15 mg/kg: time: F_(26, 386)_ = 2.197, p<0.001, knockdown: F_(1, 386)_ = 5.163, p=0.02; 30 mg/kg: time: F_(26, 339)_ = 3.569, p<0.0001, knockdown: F_(1, 339)_ = 6.226, p=0.013), and total ambulation was significantly increased as compared to controls at the 5 mg/kg (p=0.01) and 15 mg/kg (p=0.04) doses. **(d-f)** *Female locomotor activity*. Rit2 knockdown in DANs significantly decreased female response to cocaine at 15 mg/kg (time: F_(26, 333)_ = 1.702, p=0.019, knockdown: F_(1, 333)_ = 26.08, p<0.0001), and significantly decreased total ambulation in response to cocaine at the 15 mg/kg dose (p=0.04).

### DAergic Rit2 knockdown differentially impacts DAergic characteristics in males and females

The observed changes in DA-dependent behaviors suggest that DAergic Rit2 expression may impact DAergic function. Rit2 binds directly to DAT, and Rit2 activity is required for PKC-stimulated DAT endocytosis in cell lines [16]. However, it is unknown whether Rit2 impacts critical determinants of the DAergic phenotype *in vivo*, such as TH and DAT. Therefore, we measured TH and DAT mRNA levels in ventral midbrain, and their respective protein levels in *ex vivo* striatal slices, in both males and females, following injection with AAV9-TRE-shRit2 and maintenance ±dox. DAergic Rit2-KD had no significant effect on either TH or DAT mRNA levels in females (Fig. 4a,b). However, in males, Rit2-KD significantly decreased DAT and TH mRNA levels (Fig. 4c,d). DAergic Rit2-KD did not significantly affect either total striatal TH or DAT in females (Fig. 4e,f), and likewise did not affect male TH levels (Fig. 4g), despite decreased male TH mRNA levels. However, Rit2-KD significantly decreased total DAT protein in male striatum (Fig. 4h). Remarkably, decreased total male DAT was accompanied by significantly increased striatal DAT function, as measured by [^3^H]DA uptake (Fig. 4i). These surprising results prompted us to ask whether, Rit2-KD may have shifted the DAT distribution to the plasma membrane in males as a compensatory mechanism. *Ex vivo* slice biotinylation studies revealed that Rit2-KD did not significantly alter the fraction of DAT on the plasma membrane in either DS (p>0.99). or VS (p>0.99), as compared to control (eGFP) slices (Fig. 4j). Coupled with the decrease in total DAT, these results suggest that the absolute DAT level at the plasma membrane is reduced in males. Morevoer, since Rit2 plays a role in maintaining DAT protein levels in males, but not females, it is possible that these differences may have contributed to the sex-specific, differential impact on acute cocaine responses. Interestingly, baseline DAT surface distribution in controls was significantly lower in DS than VS, which suggests that mechanisms controlling DAT’s presentation at the plasma membrane may be subject to region-specific control.

**Figure 4.**
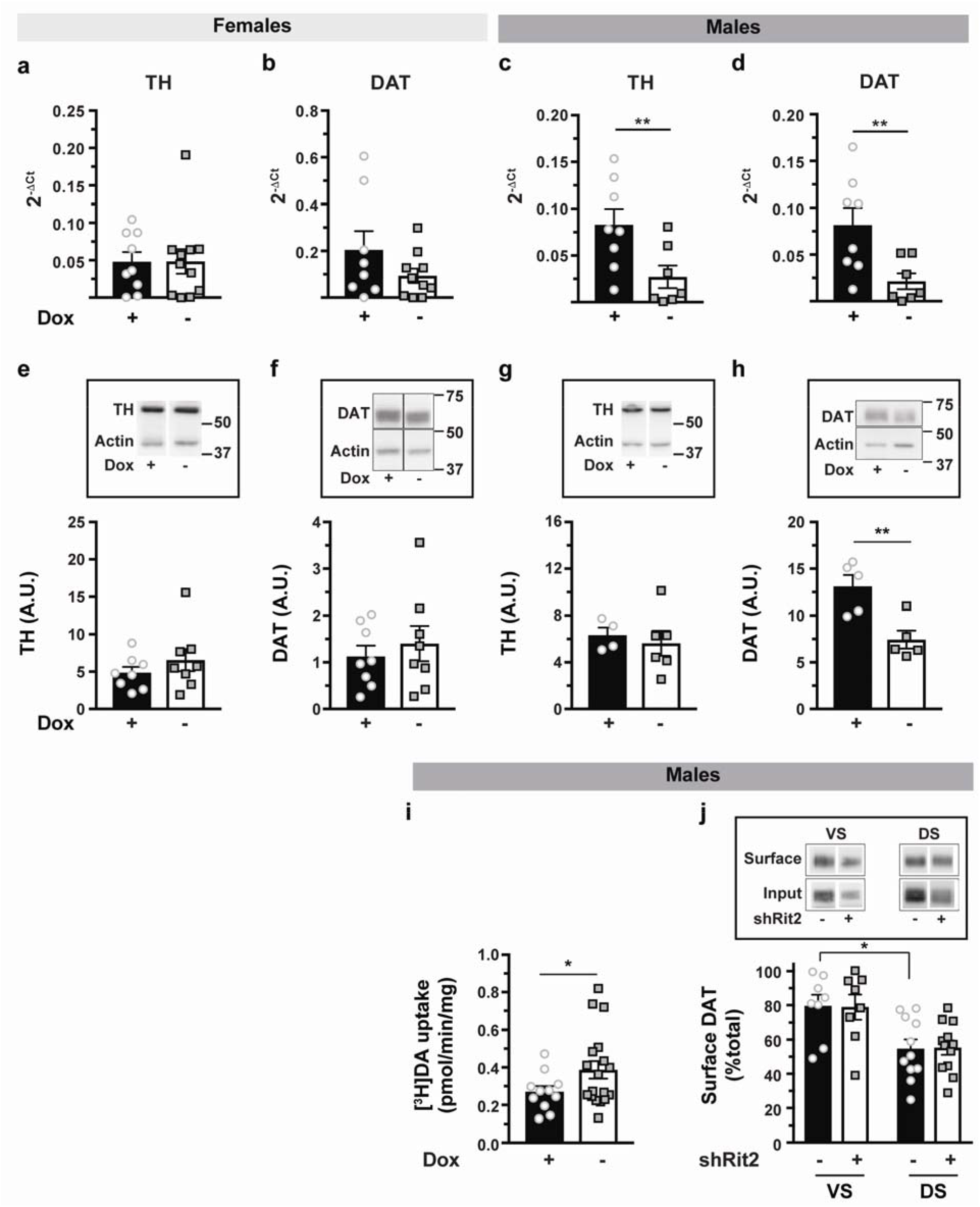
Sex-specific effects on DAergic determinants following Rit2-KD. *Pitx3^IRES2-tTA^*/+ mouse VTA were bilaterally injected either: with AAV9-TRE-shRit2 and maintained ±dox diet (a-i), or AAV9-TRE-shRit2 vs. AAV9-TRE-eGFP (j). Mice were assessed 4-6 weeks post-injection. **(a-d)** *RT-qPCR studies*. Tissue was isolated by laser capture microscopy (SNC; females) or midbrain punch (males), and the indicated mRNA levels were measured as described in *Methods*. Rit2-KD did not affect either TH (a; p=0.87) or DAT (b, p=0.52) expression in female SNc, but significantly decreased both TH (c, p=0.02) and DAT (d, p=0.014) expression in male midbrain. Asterisks indicate a significant difference from +dox control, two-tailed Student’s t test, n=7-8. **(e-h)** *Striatal immunoblots for TH and DAT*. Female (e,f) and male (g,h) striata were harvested and total striatal TH and DAT content were measured by immunoblot, as indicated. *Tops:* Representative immunoblots for the indicated proteins. *Bottoms:* Quantified band intensities, normalized to actin loading controls. **Significantly less than +dox control, p=0.006, two-tailed Student’s t test, n=5 independent mice/condition. **(i)** *[^3^H]DA uptake in ex vivo striatal slices:* DA uptake was measured in male striatal slices, as described in *Methods*. Data are presented as specific DA uptake, normalized to TH content in each slice. ‘Significantly larger than +dox control, p=0.047, two-tailed Student’s t test with Welch’s correction, n=11-18 slices from 4-6 independent mice/condition. **(j)** *Ex vivo striatal slice biotinylation*. Steady state DAT surface levels were determined in dorsal (DS) and ventral (VS) striatal slices as described in *Methods. Top:* Representative immunoblots of DAT surface and input fractions. *Bottom:* Average DAT surface distribution, presented as %total input DAT on cell surface. In control mice (-shRit2), baseline DAT surface distribution was significantly lower in dorsal vs. ventral striatum *p=0.013, one-way ANOVA with Sidak’s multiple comparison test, n=8-12 independent slices from 3-4 mice/condition. Bands displayed within each panel are from the same exposure of the same immunoblot, and were cropped for presentation purposes only.

### DAergic Rit2-KD impacts spontaneous glutamatergic input onto NAcc MSNs in a sex- and cell type-specific manner

DA in the NAcc 1) influences MSN responses to excitatory input [24], and 2) modulates the spontaneous activity of glutamatergic afferents that synapse onto MSNs [25]. Moreover, cocaine impacts many of these parameters [26,27]. Since DAergic Rit2-KD impacted both acute cocaine responses and presynaptic DAT, we hypothesized that that NAcc MSN spontaneous activity, intrinsic biophysical properties, and/or response to cocaine may be affected by DAergic Rit2-KD. To test these possibilities, we recorded from D1+ and D2+ MSNs in NAcc core, from slices prepared from male and female *Pitx3^IRES2-tTA^;D1-tdTomato* mice injected with either AAV9-TRE-eGFP or AAV9-TRE-shRit2. In males, DAergic Rit2-KD significantly decreased baseline spontaneous sEPSP (sEPSP) frequency in D1+, but not D2+, MSNs (Fig. 5a,b). In contrast, in females, DAergic Rit2-KD significantly decreased baseline sEPSP frequency in D2+, but not D1+, MSNs (Fig. 5c,d). Rit2-KD additionally decreased the input resistance in D1+, but not D2+, MSNs in both males and females (Fig. 5e-h). sEPSP amplitudes were also differentially impacted by DAergic Rit2-KD in a cell-type and sex-specific manner: In males, D1+ amplitudes were increased, while D2+ amplitudes were decreased (Fig. S4a, b), whereas in females sEPSP amplitude was increased in D1+, but unchanged in D2+, MSNs (Fig. S4c,d). In male controls, cocaine significantly decreased sEPSP frequency in D1+, but not D2+, MSNs, and DAergic Rit2-KD abolished the effect of cocaine on sEPSP frequency (Fig. 5i,j). Rit2-KD did not affect either RMP, AP threshold, AHP, or AP overshoot in either D1+ or D2+ MSNs from either sex (Fig. S4e-t). Taken together, these results demonstrate that DAergic Rit2 expression significantly impacts multiple facets of striatal physiology, with distinct, sex-specific differences.

**Figure 5.**
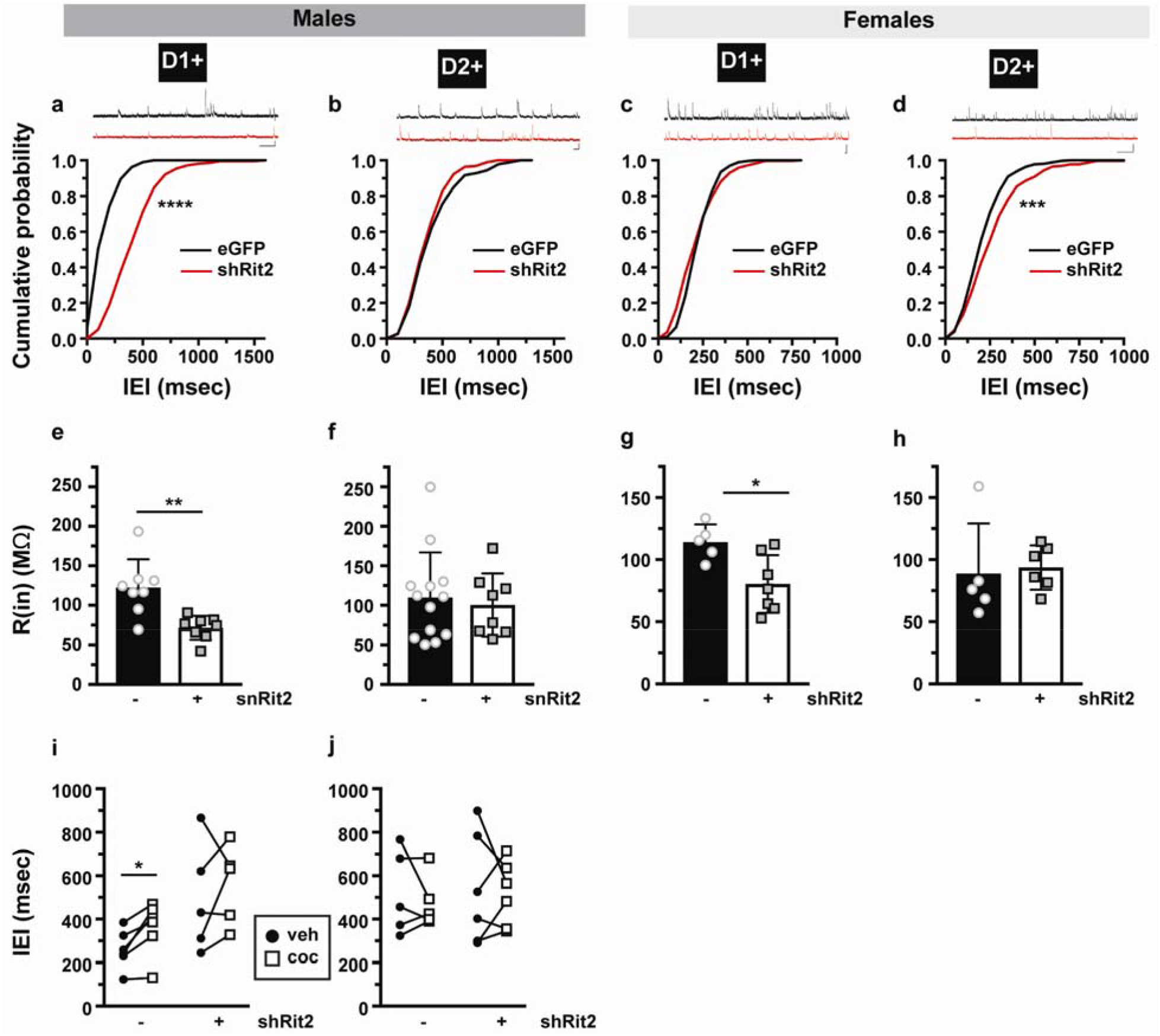
Rit2-KD in DA neurons reduces the excitatory input onto NAcc MSNs in a cell type and sex-specific manner. *Ex vivo striatal slice recordings. Pitx3^IRES-tTA^/Drd1a-tdTomato* mice were bilaterally injected with either AAV9-TRE-eGFP or AAV9-TRE-shRit2, and MSN activity was recorded in current clamp, ±10μM cocaine, 4 weeks post-injection, as described in *Methods*. Data shown are from 6-8 cells/condition recorded from 3-4 independent male and female mice each. **(a-d)** Cumulative distributions of sEPSP frequencies (interevent interval; IEI) from males and females. Representative traces are shown, above. Scale bar = 0.5 sec and 1.0mV. Asterisks indicate significant differences between eGFP vs. shRit2 cumulative distributions, two-tailed Kolmogorov-Smirnov test, ***p=0.0003, ****p<0.0001. **(e-h)** Input resistance. DAergic Rit2-KD decreased input resistance (R(in)) in D1+, but not D2+ MSNs from both males and females. Asterisks indicate significant differences between eGFP vs. shRit2 values, unpaired, two-tailed Student’s t test, *p=0.017, **p=0.002 **(I,j)** Effect of cocaine on sEPSP frequency. Cocaine treatment significantly decreased sEPSP frequenciy in D1+ cells from control (eGFP) but not shRit2 male mice. *p=0.017, paired, two-tailed Students t test.

## Discussion

DAergic transmission drives locomotor behavior both in rodents [28,29], and in humans, where DAN death underlies the movement anomalies in PD [30]. Aberrant DA signaling is a hallmark of multiple neuropsychiatric disorders, most notably addiction and schizophrenia [31,32]. Although optogenetic and chemogenetic tools, in combination with Cre/Lox technology, have advanced our understanding of the circuitry and molecular mechanisms that impact DAergic function, AAV-mediated conditional knockdown in DANs has been significantly more challenging to accomplish. Here, we demonstrated that the Tet-OFF approach can be leveraged to achieve AAV-mediated, conditional, inducible, *in vivo* gene silencing. Using this approach, we identified sex-specific roles for DAergic Rit2 expression in DA-dependent behaviors, presynaptic DAT function, and striatal physiology. We should note that GFP reporter expression was visibly lower in *Pitx3^IRES2-tTA^* SNc, as compared to the robust VTA signal. This could reflect limited viral spread away from the VTA; however the AAV9 serotype is noted for its widespread infection [33]. Pitx3 gene expression is ~6-fold higher in VTA than SNc [34]; thus is it more likely that differential Pitx3 promoter activity drove differential GFP expression in VTA vs. SNc. Nevertheless, shRit2 significantly decreased Rit2 expression in SNc, suggesting that the SNc Pitx3 promoter is sufficiently active to drive tTA, and TRE-shRit2, expression. We also observed some sparse GFP+ cells dorsomedial to the VTA, likely the periacqueductal gray. These cells were TH^−^, consistent with a previous report in *Pitx3^Cre^* mice [35]. Both *DAT^tTA^* and *TH^tTA^* mice have also been reported [36,37]. However, DAT expression in *DAT^tTA^* mice is markedly reduced, due to the tTA placement in the DAT 5’ UTR [37], and *TH^tTA^* transgenic mice exhibit incomplete tTA expression throughout the SNc and VTA [36]. Moreover, since noradrenergic neurons are TH^+^, tTA expression is not limited to DAergic neurons in *TH^tTA^* mice. Thus, *Pitx3^IRES2-tTA^* mice are currently the best option for DAergic tTA expression. Another considerable advantage to TET-OFF approach for shRNA expression is the ability to induce/suppress knockdown with doxycycline. It should be noted, however, that we occasionally observed GFP signals in TH+ cells in tissue harvested from dox-treated mice (not shown). This is likely the reason that Rit2 mRNA levels were significantly more variable in (+)dox mice vs. (−)dox mice (Fig. 1h,i), and may be attributed to either the short dox half-life, and/or the inability to strictly control mouse dox dosing via *ad libitum* route. Nevertheless, given the robust *in vivo* knockdown and behavioral phenotypes we observed, we believe that the tTA/TRE approach for gene silencing will have broad utility in future longitudinal behavioral/addiction studies, where gene silencing could be temporally controlled across varying stages of establishing, extinguishing and reinstating psychostimulant reward.

Rit2 expression increases throughout postnatal development and stabilizes 3-4 weeks postnatally [9,38]. To assure that Rit2-KD in our studies did not confound the Rit2 developmental timecourse, we assessed Rit2 mRNA expression in VTA-enriched tissue in wildtype mice at 4, 7, and 11 weeks, and observed no significant differences in Rit2 expression within this timeframe (Fig. S5). Thus it is unlikely that Rit2-KD performed during this window interfered with any Rit2-dependent developmental mechanisms. Rit2 is required for neurotrophic signaling and constitutively active Rit2 expression results in neurite formation in PC6 cells [10]. Thus, it is possible that the observed changes in DA-dependent behaviors and MSN physiology following Rit2-KD may reflect a trophic impact on DA neuron viability. Rit2-KD did not affect either TH or DAT gene expression in females, but did significantly decrease TH and DAT mRNA in males (Fig. 4c,d), which was accompanied decreased DAT, but not TH, striatal protein (Fig. 4g,h). Given that we did not detect any motor deficits in either males or females in the OFT (Fig. 2), it is unlikely that DAT and TH mRNA losses reflect diminished DA neuron viability. Indeed, global Rit2 knockout did not affect neuronal development [39]. Thus, it is more likely that DAergic Rit2 loss impacted intrinsic DAergic function, rather than DA neuron viability. Future studies probing the effectof DAergic Rit2-KD on spontaneous and evoked DA release kinetics, as well as post-synaptic DA signaling should be illuminating in this regard.

Recent GWAS studies reported Rit2 as a risk allele for several DA-associated disorders, including PD, ET, ASD, schizophrenia, and bipolar disorder [14]. The link between Rit2 expression and schizophrenia is particularly interesting. A recent report examining genome-wide human 5’ short tandem repeats (STRs) found that the RIT2 locus has among the longest GA-STRs in humans, at 11 repeats, and that increased STR length drives increased Rit2 expression [14]. Moreover, a 5/5 STR Rit2 variant was identified in a proband with profound early onset psychoses, raising the possibility that decreased Rit2 expression may be linked to schizophrenia. This is further supported by a previous report that identified increased RIT2 CNVs in schizophrenic populations that result in loss of Rit2 expression [15]. Although these Rit2 perturbations would be expected at germline throughout the CNS, acute Rit2-KD in the adult animal may have potential to recapitulate DAergic dysfunction inherent in these disorders.

DAergic Rit2 silencing significantly reduced baseline anxiety, but did not affect baseline locomotor activity (Fig. 2). Mounting evidence suggests that DA signaling is involved in rodent anxiety-like behaviors. NMDA-mediated phasic bursting is required to elicit anxiety in response to aversive conditioning [40], and optogenetic studies demonstrated that activating the VTA➔PFC projection is sufficient to generate anxiety and place aversion [23]. Moreover, increased DA signaling and reduced glutamatergic firing within the NAcc is associated with increased anxiety [2]. A link between reduced NAcc DAT expression and anxiolytic behavior was recently reported [41], also consistent with our findings. Given that we used an AAV9 viral serotype, Rit2-KD likely occurred across multiple DAergic circuits, including VTA➔PFC, VTA➔NAc, and SNc➔DS. Therefore we cannot distinguish whether Rit2-KD in a specific DAergic subpopulation mediates the anxiolytic phenotype. Additionally, DANs are intrinsically heterogeneous [42], and recent studies further suggest that both DA and glutamate are released from independent vesicle populations within DAergic neurons [43]. Thus, it is possible specific DAergic subpopulations may be more sensitive to Rit2-KD than others. It is also possible that Rit2-KD impacts intrinsic DA neuron excitability, and thereby alters DAergic response to anxiety-related inputs in the somatodendritic regions [40]. Future studies evaluating how Rit2 expression impacts DAN physiological properties should shed further light on these possibilities.

DAergic Rit2-KD exerted striking, sex-specific effects on acute cocaine responses. In males, Rit2-KD increasd acute cocaine sensitivity (Fig. 3). Previous studies reported increased cocaine-induced hyperactivity in male DAT-KD mice [44], and that cocaine-hypersensitive male rats express less DAT [45]. These results raise the possibility that the enhanced cocaine sensitivity we observed in Rit2-KD males might be due to their decreased DAT levels (Fig 4d). In contrast, in females, Rit2-KD abolished hyperlocomotion at 15 mg/kg cocaine (Fig. 3b), whereas DAT protein levels were unaffected (Fig. 4f). Females exhibit more robust addictive behaviors than males [46], and female mice in estrous are more sensitive to cocaine’s rewarding effects and have increased DAT N-terminal phosphorylation, as compared to dioestrous females [47]. Thus, hormonal signaling in females may contribute both to basal increased cocaine sensitivity, and to loss of cocaine sensitivity in response to Rit2-KD. Ongoing studies will further illuminate potential sex differences in DAT endocytic mechanisms, and whether these might contribute directly to the Rit2-KD phenotype. Interestingly, the Nestler laboratory reported that chronic cocaine treatment increased ÁFosB binding to the Rit2 promotor in mouse NAcc [48], suggesting that Rit2 may play a wider role within the reward circuitry, beyond its role in DANs.

In addition to imposing sex-specific effects on acute cocaine sensitivity, DAergic Rit2-KD also significantly altered the excitatory drive onto NAcc MSNs in a cell type- and sex-specific manner. sEPSP frequency was decreased in male D1+ MSNs, whereas in females it was decreased in female D2+ MSN populations. D1+ MSN activation is canonically linked to motivation and reward, whereas D2+ MSN activation is thought to mediate drug aversion [49,50]. However, recent studies challenge this dogma, and provide compelling evidence that, unlike DS MSNS, NAcc MSN D1+ and D2+ outputs are significantly more complex than had been previously appreciated [51]. Moreover, a recent optogenetic study directly demonstrated that NAcc D2+ MSN activation drives a positive motivational valence [52]. Thus, the differential effect of DAergic Rit2-KD on D1+ and D2+ excitatory drive in males and females may potentially underlie the reciprocal changes in cocaine sensitivity we observed following Rit2-KD. Cocaine-induced decreases in sEPSP frequency have been reported *ex vivo* in NAcc [53], and *in vivo* in PFC [54], and requires presynaptic D1R activation on glutamatergic terminals in *ex vivo* NAcc [53]. We likewise found that cocaine significantly decreased sEPSP frequency in control male D1+, but not D2+, MSNs, and DAergic Rit2-KD abolished cocaine-induced decreases in sEPSP frequency (Fig. 5i,j). The dampened cocaine effect on D1+ MSNs could be directly due to decreased DAT (Fig. 4h), or could be due to a floor effect on glutamatergic sEPSPs, since Rit2-KD itself decreased sEPSP frequency in D1+ MSNs (Fig. 5a). Interestingly, similar changes in MSN sEPSPs were reported in DAT-KD mice, which express 90% less DAT than wildtype mice [55]. We also found that DAergic Rit2-KD significantly decreased D1+, but not D2+, MSN input resistance in both males and females (Fig. 5e-h). DA has broad effects on MSN excitability, both in the direct and indirect basal ganglia output neurons [56]. Thus, any changes in DA signaling would be expected to influence MSN excitability and are consistent with our findings.

DAT expression and function have multimodal effects on DAergic homeostasis and DA-dependent behaviors. Presynaptic DA content is highly sensitive to DAT expression and function, where DA recapture is required to maintain DAergic tone [7]. Given that we found that DAergic Rit2-KD decreased male total striatal DAT protein (Fig. 4h), it is possible that DAT losses compromised DA signaling, which downstream impacted MSN function and cocaine responses. Paradoxically, despite losses in DAT protein, we measured significantly more DA uptake in striatal slices following Rit2-KD (Fig. 4i). It should be noted that D2R activation can drive DAT plasma membrane insertion [57,58]. Taken together with our previous findings that Rit2 is required for regulated DAT endocytosis [16], it is possible that in the course of measuring DA uptake, D2R were activated and drove DAT insertion. In Rit2-KD slices, DAT may have not been effectively retrieved from the plasma membrane, yielding a net increase in uptake. Since we did not include D2R antagonists in our DA uptake studies, this is entirely possible. Future studies measuring DA release and clearance, and how regulated DAT trafficking mechanisms impact this process, will directly test these possibilities. It is additionally noteworthy that the DAT baseline distribution between plasma membrane and endocytic vesicles was significantly lower in DS than VS, suggesting that there are region-specific differences in DAT endocytic trafficking. Similar results were recently reported for a knock-in mouse expressing the ADHD-associated DAT coding variant, A559V, in which DAT A559V basal distribution was significantly lower in DS than VS [58].

Taken as a whole, these studies are the first to demonstrate a role for Rit2 GTPase in DA-related behaviors and striatal physiology. Morevoer, the approach we developed for conditional, inducible gene silencing in DANs is likey to facilitate future studies examining DAergic mechanisms involved in DA-related neuropsychiatric disorders.

## Acknowlegements

The authors would like to acknowledge Dr. Susanna Casacuberta for outstanding surgical training, and Tucker Conklin for excellent technical assistance.

## References

1 Wise RA. Dopamine, learning and motivation. Nat Rev Neurosci. 2004;5(6):483–94.

2 Russo SJ, Nestler EJ. The brain reward circuitry in mood disorders. Nat Rev Neurosci. 2013;14(9):609–25.

3 Calhoon GG, Tye KM. Resolving the neural circuits of anxiety. Nat Neurosci. 2015;18(10):1394–404.

4 Poewe W, Seppi K, Tanner CM, Halliday GM, Brundin P, Volkmann J, et al. Parkinson disease. Nature reviews Disease primers. 2017;3:17013.

5 Faraone SV, Asherson P, Banaschewski T, Biederman J, Buitelaar JK, Ramos-Quiroga JA, et al. Attention-deficit/hyperactivity disorder. Nature reviews Disease primers. 2015;1:15020.

6 Grace AA. Dysregulation of the dopamine system in the pathophysiology of schizophrenia and depression. Nat Rev Neurosci. 2016;17(8):524–32.

7 Kristensen AS, Andersen J, Jorgensen TN, Sorensen L, Eriksen J, Loland CJ, et al. SLC6 neurotransmitter transporters: structure, function, and regulation. Pharmacological reviews. 2011;63(3):585–640.

8 Zhou Q, Li J, Wang H, Yin Y, Zhou J. Identification of nigral dopaminergic neuron-enriched genes in adult rats. Neurobiol Aging. 2011;32(2):313–26.

9 Spencer ML, Shao H, Tucker HM, Andres DA. Nerve growth factor-dependent activation of the small GTPase Rin. J Biol Chem. 2002;277(20):17605–15.

10 Shi GX, Han J, Andres DA. Rin GTPase couples nerve growth factor signaling to p38 and b-Raf/ERK pathways to promote neuronal differentiation. J Biol Chem. 2005;280(45):37599–609.

11 Uenaka T, Satake W, Cha PC, Hayakawa H, Baba K, Jiang S, et al. In silico drug screening by using genome-wide association study data repurposed dabrafenib, an antimelanoma drug, for Parkinson’s disease. Hum Mol Genet. 2018;27(22):3974–85.

12 Hamedani SY, Gharesouran J, Noroozi R, Sayad A, Omrani MD, Mir A, et al. Ras-like without CAAX 2 (RIT2): a susceptibility gene for autism spectrum disorder. Metab Brain Dis. 2017;32(3):751–55.

13 Daneshmandpour Y, Darvish H, Emamalizadeh B. RIT2: responsible and susceptible gene for neurological and psychiatric disorders. Mol Genet Genomics. 2018;293(4):785–92.

14 Emamalizadeh B, Jamshidi J, Movafagh A, Ohadi M, Khaniani MS, Kazeminasab S, et al. RIT2 Polymorphisms: Is There a Differential Association? Mol Neurobiol. 2017;54(3):2234–40.

15 Glessner JT, Reilly MP, Kim CE, Takahashi N, Albano A, Hou C, et al. Strong synaptic transmission impact by copy number variations in schizophrenia. Proc Natl Acad Sci U S A. 2010;107(23):10584–9.

16 Navaroli DM, Stevens ZH, Uzelac Z, Gabriel L, King MJ, Lifshitz LM, et al. The plasma membrane-associated GTPase Rin interacts with the dopamine transporter and is required for protein kinase C-regulated dopamine transporter trafficking. J Neurosci. 2011;31(39):13758–70.

17 Xie J, Mao Q, Tai PWL, He R, Ai J, Su Q, et al. Short DNA Hairpins Compromise Recombinant Adeno-Associated Virus Genome Homogeneity. Mol Ther. 2017;25(6):1363–74.

18 Wu Z, Yang H, Colosi P. Effect of genome size on AAV vector packaging. Mol Ther. 2010;18(1):80–6.

19 Gabriel LR, Wu S, Kearney P, Bellve KD, Standley C, Fogarty KE, et al. Dopamine transporter endocytic trafficking in striatal dopaminergic neurons: differential dependence on dynamin and the actin cytoskeleton. J Neurosci. 2013;33(45):17836–46.

20 Wu S, Bellve KD, Fogarty KE, Melikian HE. Ack1 is a dopamine transporter endocytic brake that rescues a trafficking-dysregulated ADHD coding variant. Proc Natl Acad Sci U S A. 2015;112(50):15480–5.

21 Lin X, Parisiadou L, Sgobio C, Liu G, Yu J, Sun L, et al. Conditional expression of Parkinson’s disease-related mutant alpha-synuclein in the midbrain dopaminergic neurons causes progressive neurodegeneration and degradation of transcription factor nuclear receptor related 1. J Neurosci. 2012;32(27):9248–64.

22 Beninger RJ. The role of dopamine in locomotor activity and learning. Brain research. 1983;287(2):173–96.

23 Gunaydin LA, Grosenick L, Finkelstein JC, Kauvar IV, Fenno LE, Adhikari A, et al. Natural neural projection dynamics underlying social behavior. Cell. 2014;157(7):1535–51.

24 Swapna I, Bondy B, Morikawa H. Differential Dopamine Regulation of Ca(2+) Signaling and Its Timing Dependence in the Nucleus Accumbens. Cell Rep. 2016;15(3):563–73.

25 Nicola SM, Malenka RC. Dopamine depresses excitatory and inhibitory synaptic transmission by distinct mechanisms in the nucleus accumbens. J Neurosci. 1997;17(15):5697–710.

26 Cepeda C, Buchwald NA, Levine MS. Neuromodulatory actions of dopamine in the neostriatum are dependent upon the excitatory amino acid receptor subtypes activated. Proc Natl Acad Sci U S A. 1993;90(20):9576–80.

27 Nicola SM, Malenka RC. Modulation of synaptic transmission by dopamine and norepinephrine in ventral but not dorsal striatum. Journal of neurophysiology. 1998;79(4):1768–76.

28 Marshall JF, Berrios N. Movement disorders of aged rats: reversal by dopamine receptor stimulation. Science. 1979;206(4417):477–9.

29 Starr BS, Starr MS. Differential effects of dopamine D1 and D2 agonists and antagonists on velocity of movement, rearing and grooming in the mouse. Implications for the roles of D1 and D2 receptors. Neuropharmacology. 1986;25(5):455–63.

30 Lotharius J, Brundin P. Pathogenesis of Parkinson’s disease: dopamine, vesicles and alpha-synuclein. Nat Rev Neurosci. 2002;3(12):932–42.

31 Howes O, McCutcheon R, Stone J. Glutamate and dopamine in schizophrenia: an update for the 21st century. J Psychopharmacol. 2015;29(2):97–115.

32 Volkow ND, Wise RA, Baler R. The dopamine motive system: implications for drug and food addiction. Nat Rev Neurosci. 2017;18(12):741–52.

33 Aschauer DF, Kreuz S, Rumpel S. Analysis of transduction efficiency, tropism and axonal transport of AAV serotypes 1, 2, 5, 6, 8 and 9 in the mouse brain. PLoS One. 2013;8(9):e76310.

34 Korotkova TM, Ponomarenko AA, Haas HL, Sergeeva OA. Differential expression of the homeobox gene Pitx3 in midbrain dopaminergic neurons. The European journal of neuroscience. 2005;22(6):1287–93.

35 Smidt MP, von Oerthel L, Hoekstra EJ, Schellevis RD, Hoekman MF. Spatial and temporal lineage analysis of a Pitx3-driven Cre-recombinase knock-in mouse model. PLoS One. 2012;7(8):e42641.

36 Tillack K, Aboutalebi H, Kramer ER. An Efficient and Versatile System for Visualization and Genetic Modification of Dopaminergic Neurons in Transgenic Mice. PLoS One. 2015;10(8):e0136203.

37 Cagniard B, Beeler JA, Britt JP, McGehee DS, Marinelli M, Zhuang X. Dopamine scales performance in the absence of new learning. Neuron. 2006;51(5):541–7.

38 Lee CH, Della NG, Chew CE, Zack DJ. Rin, a neuron-specific and calmodulin-binding small G-protein, and Rit define a novel subfamily of ras proteins. J Neurosci. 1996;16(21):6784–94.

39 Pannell M, Cai W, Brelsfoard J, Carlson S, Littlejohn E, Stewart T, et al. Rin GTPase deficiency promotes neuroprotection following traumatic brain injury. The FASEB Journal. 2015;29(1_supplement):727.15.

40 Zweifel LS, Fadok JP, Argilli E, Garelick MG, Jones GL, Dickerson TM, et al. Activation of dopamine neurons is critical for aversive conditioning and prevention of generalized anxiety. Nat Neurosci. 2011;14(5):620–6.

41 Bahi A, Dreyer J-L. Dopamine transporter (DAT) knockdown in the nucleus accumbens improves anxiety-and depression-related behaviors in adult mice. Behavioural Brain Research. 2019;359:104–15.

42 Burke DA, Rotstein HG, Alvarez VA. Striatal Local Circuitry: A New Framework for Lateral Inhibition. Neuron. 2017;96(2):267–84.

43 Silm K, Yang J, Marcott PF, Asensio CS, Eriksen J, Guthrie DA, et al. Synaptic Vesicle Recycling Pathway Determines Neurotransmitter Content and Release Properties. Neuron. 2019.

44 Tilley MR, Cagniard B, Zhuang X, Han DD, Tiao N, Gu HH. Cocaine reward and locomotion stimulation in mice with reduced dopamine transporter expression. BMC Neurosci. 2007;8:42.

45 Nelson AM, Larson GA, Zahniser NR. Low or high cocaine responding rats differ in striatal extracellular dopamine levels and dopamine transporter number. J Pharmacol Exp Ther. 2009;331(3):985–97.

46 Becker JB, Koob GF. Sex Differences in Animal Models: Focus on Addiction. Pharmacological reviews. 2016;68(2):242–63.

47 Calipari ES, Juarez B, Morel C, Walker DM, Cahill ME, Ribeiro E, et al. Dopaminergic dynamics underlying sex-specific cocaine reward. Nat Commun. 2017;8:13877.

48 Renthal W, Kumar A, Xiao G, Wilkinson M, Covington HE, 3rd, Maze I, et al. Genome-wide analysis of chromatin regulation by cocaine reveals a role for sirtuins. Neuron. 2009;62(3):335–48.

49 Lobo MK, Covington HE, 3rd, Chaudhury D, Friedman AK, Sun H, Damez-Werno D, et al. Cell type-specific loss of BDNF signaling mimics optogenetic control of cocaine reward. Science. 2010;330(6002):385–90.

50 Kravitz AV, Tye LD, Kreitzer AC. Distinct roles for direct and indirect pathway striatal neurons in reinforcement. Nat Neurosci. 2012;15(6):816–8.

51 Kupchik YM, Brown RM, Heinsbroek JA, Lobo MK, Schwartz DJ, Kalivas PW. Coding the direct/indirect pathways by D1 and D2 receptors is not valid for accumbens projections. Nat Neurosci. 2015;18(9):1230–2.

52 Soares-Cunha C, Coimbra B, David-Pereira A, Borges S, Pinto L, Costa P, et al. Activation of D2 dopamine receptor-expressing neurons in the nucleus accumbens increases motivation. Nat Commun. 2016;7:11829.

53 Nicola SM, Kombian SB, Malenka RC. Psychostimulants depress excitatory synaptic transmission in the nucleus accumbens via presynaptic D1-like dopamine receptors. J Neurosci. 1996;16(5):1591–604.

54 Trantham-Davidson H, Lavin A. Acute cocaine administration depresses cortical activity. Neuropsychopharmacology. 2004;29(11):2046–51.

55 Wu N, Cepeda C, Zhuang X, Levine MS. Altered corticostriatal neurotransmission and modulation in dopamine transporter knock-down mice. Journal of neurophysiology. 2007;98(1):423–32.

56 Gerfen CR, Surmeier DJ. Modulation of striatal projection systems by dopamine. Annu Rev Neurosci. 2011;34:441–66.

57 Chen R, Daining CP, Sun H, Fraser R, Stokes SL, Leitges M, et al. Protein kinase Cbeta is a modulator of the dopamine D2 autoreceptor-activated trafficking of the dopamine transporter. J Neurochem. 2013;125(5):663–72.

58 Gowrishankar R, Gresch PJ, Davis GL, Katamish RM, Riele JR, Stewart AM, et al. Region-Specific Regulation of Presynaptic Dopamine Homeostasis by D2 Autoreceptors Shapes the In Vivo Impact of the Neuropsychiatric Disease-Associated DAT Variant Val559. J Neurosci. 2018;38(23):5302–12.

59 Fritsch R, de Krijger I, Fritsch K, George R, Reason B, Kumar MS, et al. RAS and RHO families of GTPases directly regulate distinct phosphoinositide 3-kinase isoforms. Cell. 2013;153(5):1050–63.

60 Mueller C, Ratner D, Zhong L, Esteves-Sena M, Gao G. Production and discovery of novel recombinant adeno-associated viral vectors. Curr Protoc Microbiol. 2012;Chapter 14:Unit14D 1.

61 Burger C, Nguyen FN, Deng J, Mandel RJ. Systemic mannitol-induced hyperosmolality amplifies rAAV2-mediated striatal transduction to a greater extent than local co-infusion. Mol Ther. 2005;11(2):327–31.

62 Schmittgen TD, Livak KJ. Analyzing real-time PCR data by the comparative C(T) method. Nat Protoc. 2008;3(6):1101–8.

63 Staal RGW, Rayport S, Sulzer D. Amperometric Detection of Dopamine Exocytosis from Synaptic Terminals. In: Michael AC, Borland LM, editors. Electrochemical Methods for Neuroscience. Boca Raton (FL); 2007.

