## Supplemental Figures for "Conditional, inducible gene silencing in dopamine neurons reveals a sex-specific role for Rit2 GTPase in acute cocaine response and striatal function"

**Sweeny et al.**

**SUPPLEMENTAL FIGURES**

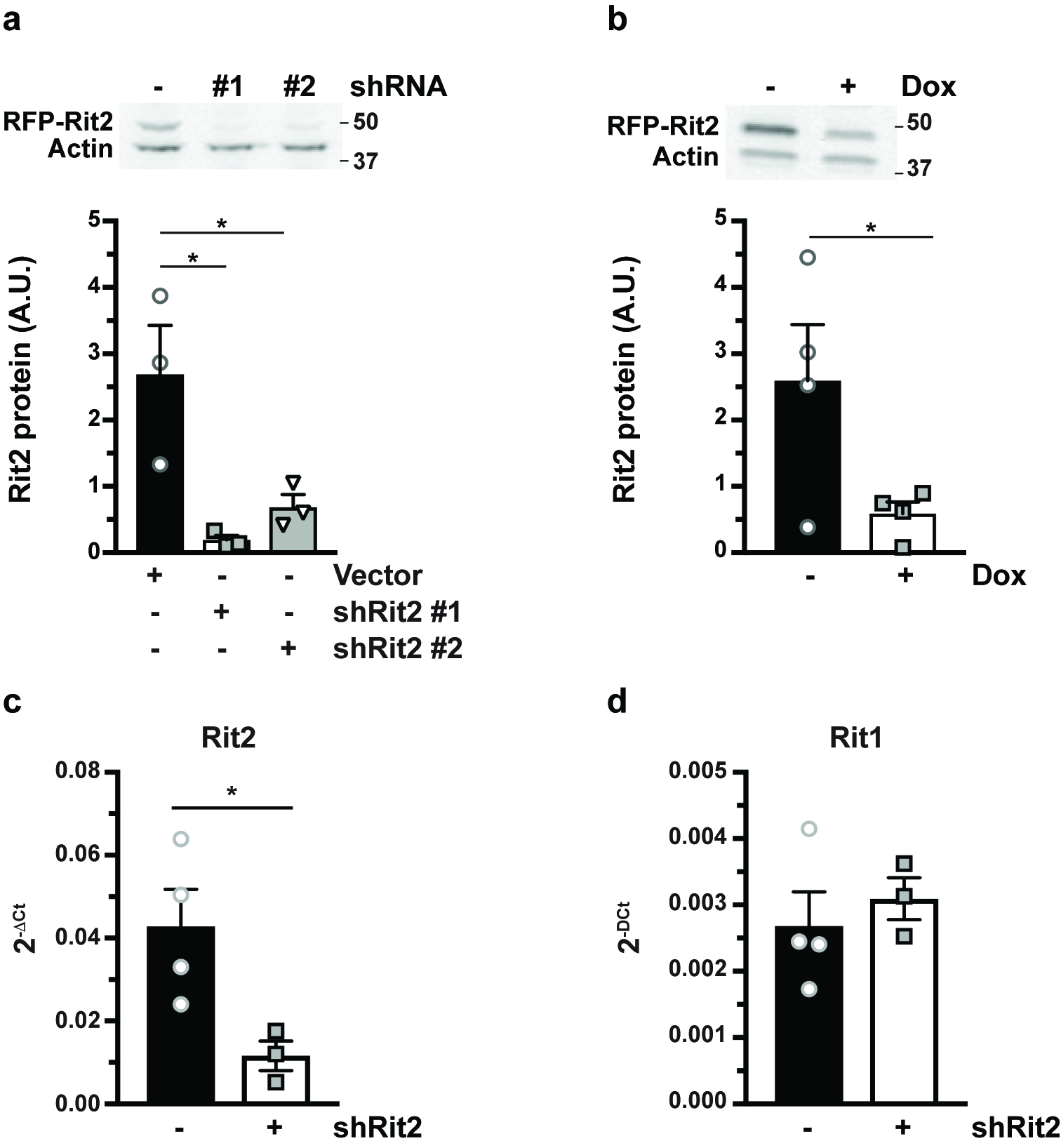

**Figure S1**. **Identification of mRit2-directed shRNAs.** **(a, b)** *In vitro Rit2 knockdown studies*. **(a)** HEK293T cells were transiently co-transfected with RFP-mRit2 reporter and either vector (pGIPZ) or the indicated Rit2-directed shRNAs cloned into the pGIPZ vector. RFP-Rit2 protein levels were assessed 48 hours post-transfection by immunoblot, normalized to actin loading controls. *Significantly different from vector control, p<0.04, one-way ANOVA with Dunnett’s multiple comparisons test, n=3. **(b)** Test for TET-inducible shRNA expression with the pAAV-TRE-shRit2 vector. Cells were co-transfected with RFP-Rit2 reporter, rtTA, and pAAV-TRE-shRit2, and were treated ±500ng/mL dox for 48 hours. *Significantly less than (-)dox control, p=0.02, one-tailed Student’s t test, n=4. **(c,d)** *In vivo shRit2 specificity studies.* 3 week old *Pitx3^IRES2-tTA^* mice VTA were bilaterally injected with AAV9-TRE-shRit2 and were maintained ±dox diet for 4-5 weeks. Ventral midbrain tissue punches were obtained and mRit2 **(c)** and mRit1 **(d)** mRNA levels were determined in parallel by RT-qPCR. AAV9-TRE-shRit2 significantly decreased mRit2 expression in (-)dox mice, as compared to (+)dox controls (p=0.02, one-tailed Student’s t test, n=3-4), but did not affect mRit1 expression (p=0.56).

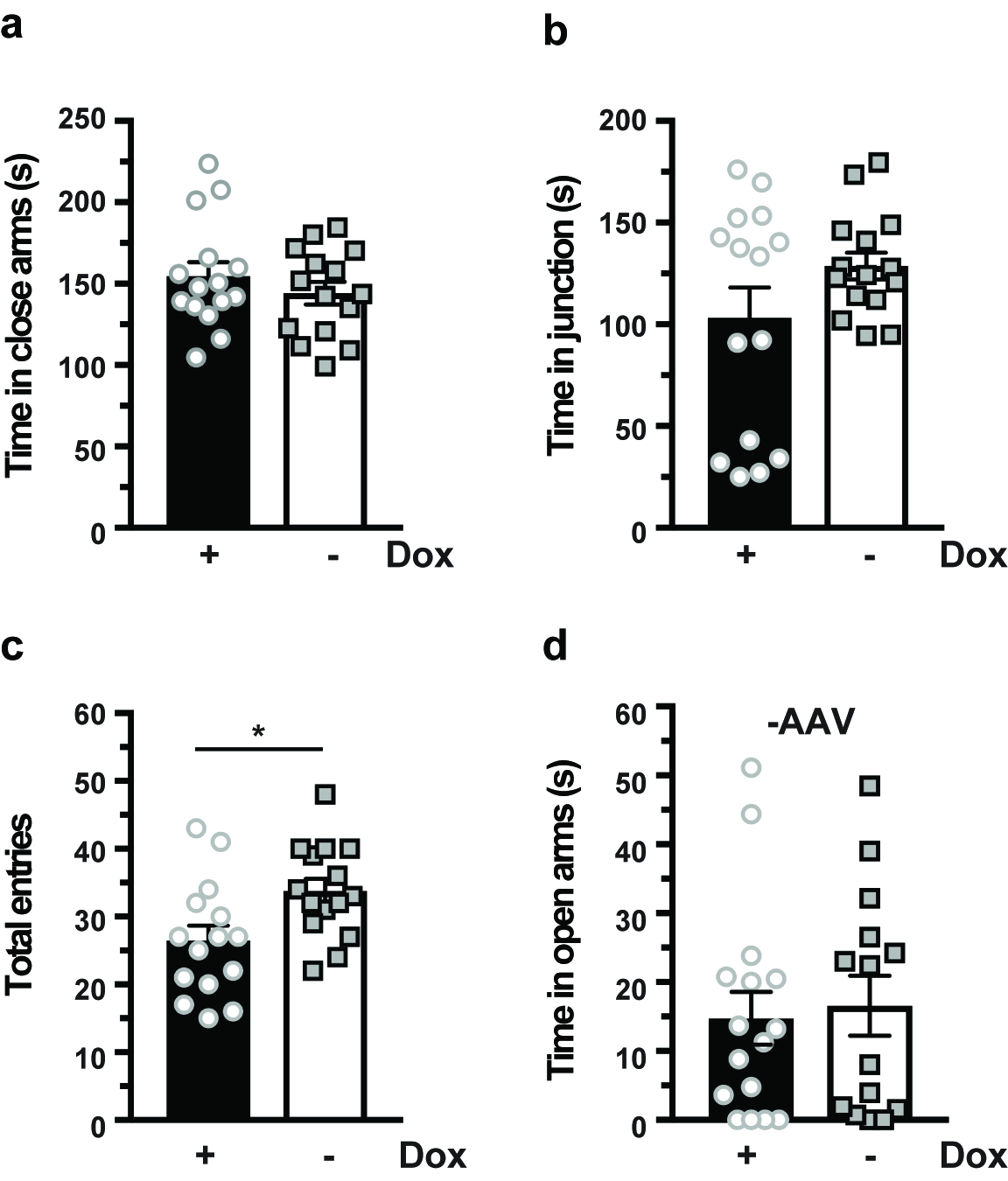

**Figure S2**. **Elevated Plus Maze.** AAV9-TRE-shRit2 did not affect time spent in the closed arms (**a;** p=0.35), or in the junction (**b;** p=0.13), in (-)dox mice, as compared to (+)dox control mice, but signficantly increased the total number of entries (**c;** *p<0.05, two-tailed Student’s t test, n=14-16). **(d)** *Effect of doxycycline diet on time spent in open arms.* Näive *Pitx3^IRES2-tTA^/+* mice were maintained ±dox diet for a minimum 4 weeks. Diet did not affect time spent in the open arms (p=0.75, two-tailed Student’s t test, n=16 pooled males and females).

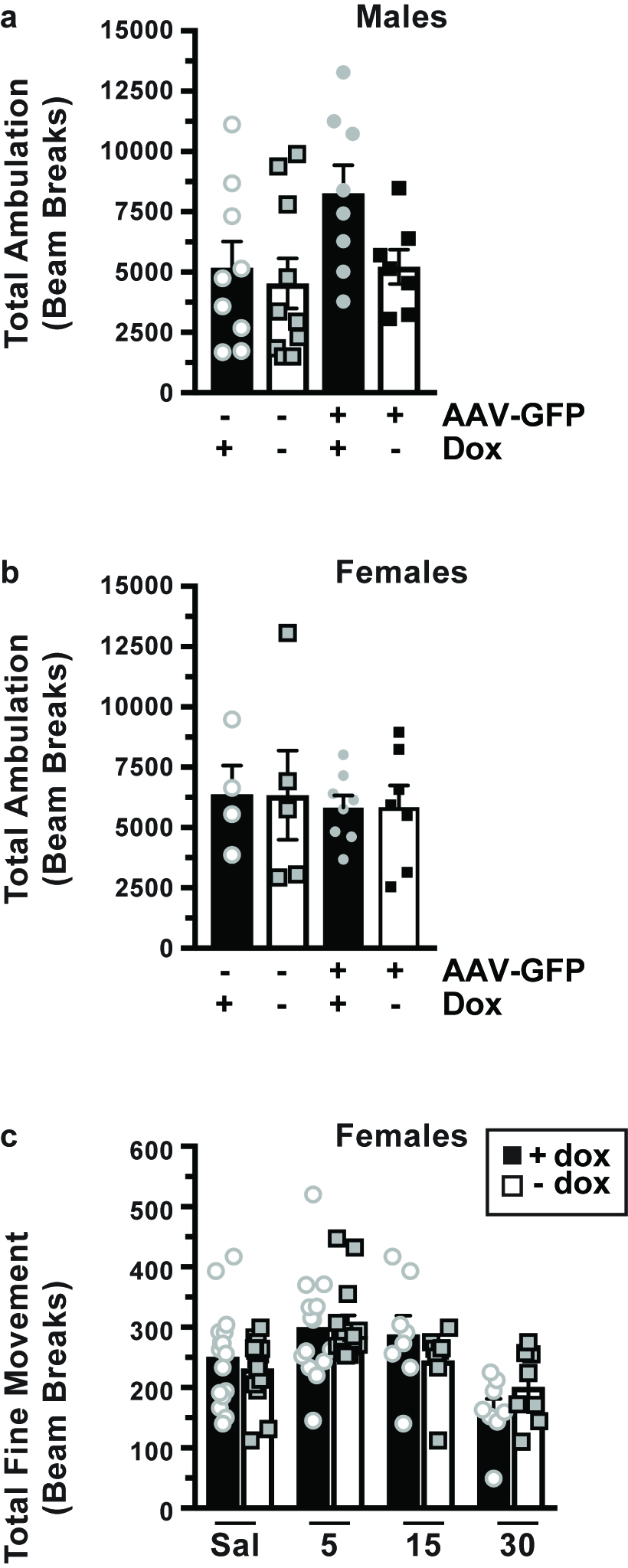

**Figure S3. Controls for acute cocaine hyperlocomotion. (a, b)** *Cocaine-induced locomotor activity.* Male and female *Pitx3^IRES2-tTA^/+* mice VTA were either bilaterally injected with AAV9-TRE-GFP, or were non-injected, and were maintained ±dox diet for 4 weeks minimum. Locomotor activity was monitored following a single, 15mg/kg I.P. cocaine injection. **(a)** *Males*: Neither viral injection nor dox diet significantly affected locomotor response to cocaine (p=0.06, one-way ANOVA, n=7-11). **(b)** *Females:* Neither viral injection nor dox diet impact locomotor response to cocaine (p=0.97, one-way ANOVA, n=4-8). **(c)** *Effect of Rit2 silencing on fine movement in females*. Female *Pitx3^IRES2-tTA^/+* mice VTA were bilaterally injected with AAV9-TRE-shRit2 were maintained ±dox diet for 6 weeks minimum. Fine movement was measured in response to I.P. injection with either saline (Sal), or 5, 15, 30 mg/kg cocaine. DAergic Rit2 knockdown had no significant effect on fine movement at any doses tested, as compared to +dox controls. Saline: p=0.43, n=16; 5mg/kg: p=0.89, n=15, 15mg/kg: p=0.26, n=8; 30mg/kg: p=0.20, n=8; upaired, two-tailed Student’s t test, n=7-15.

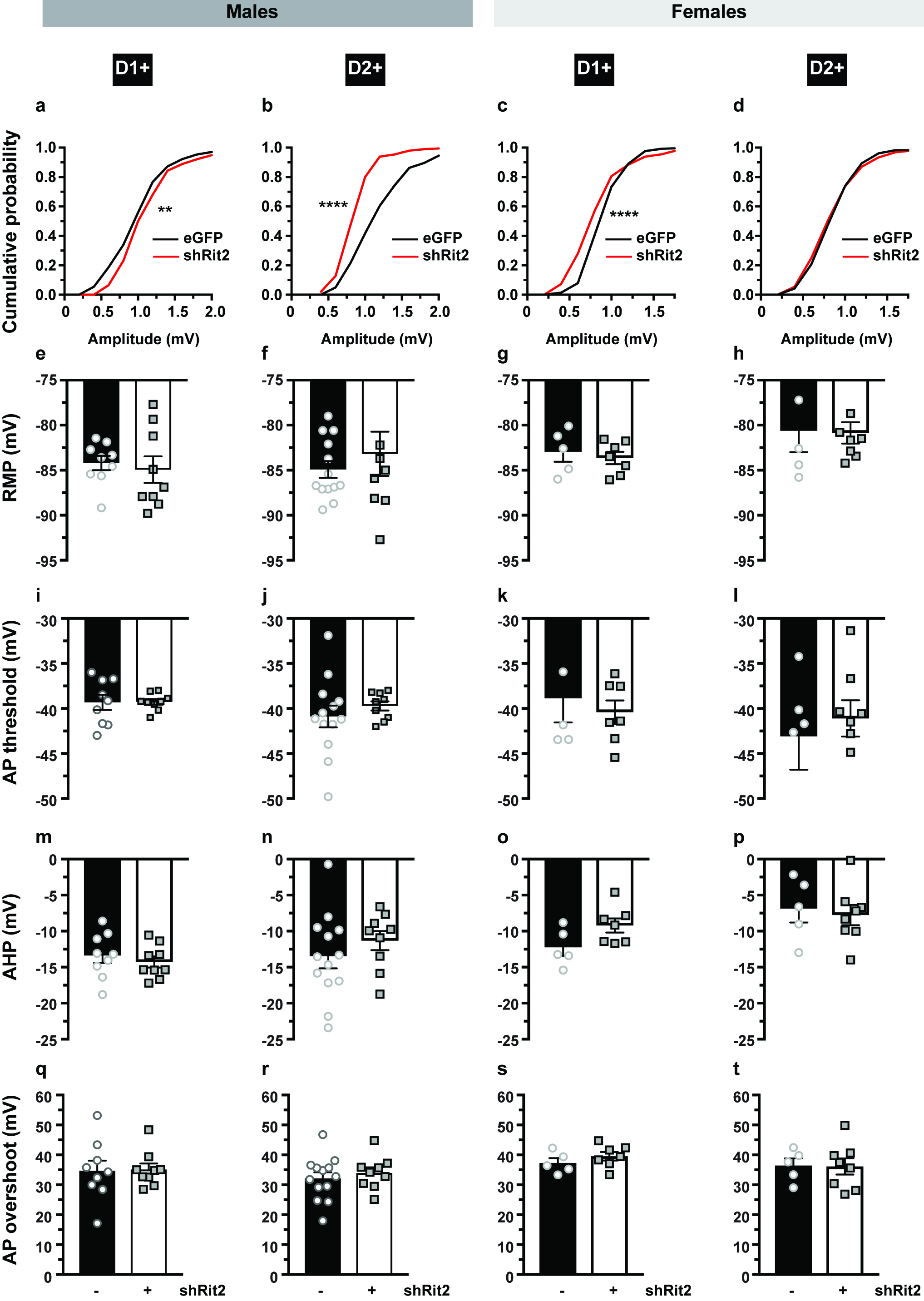

**Figure S4. Effect of DAergic Rit2-KD on sEPSP amplitudes and MSN intrinsic properties.** *Pitx3^IRES-tTA^/Drd1a-tdTomato* mice expressed AAV9-TRE-eGFP or AAV9-TRE-shRit2-eGFP for 4 weeks, and MSNs were evaluated under current clamp as described in *Methods*. **(a-d).** Cumulative distributions of sEPSP amplitudes (mV) from males and females. Asterisks indicate significant differences between eGFP vs. shRit2 cumulative distributions, two-tailed Kolmogorov-Smirnov test, **p=0.004, ****p<0.0001. **(e-t)** DAergic Rit2-KD had no significant effect on either resting membrane potential (RMP; **e-h**), action potential (AP) threshold **(i-l)**, after-hyperpolerization potential (AHP; **m-p)**, or AP overshoot **(q-t)**, as compared to MSNs from control (-shRit2) mice. Two-tailed Student’s t test.

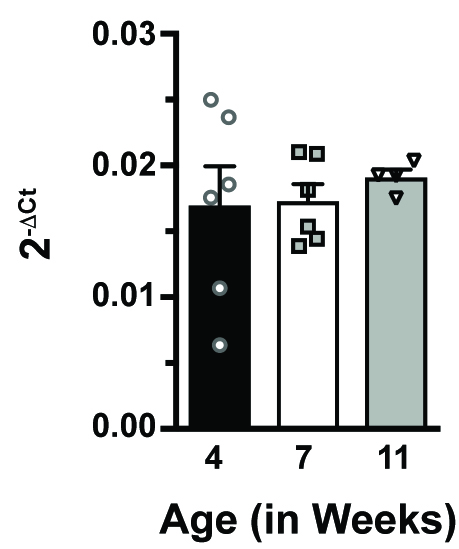

**Figure S5. Midbrain** **Rit2 mRNA expression is stable after 4 weeks of age.** Mouse midbrain was isolated from wildtype mice, and Rit2 expression was measured by RT-qPCR, normalized to GAPDH internal controls. Rit2 expression did not differ significantly across ages 4, 7, and 11 weeks of age (p=0.78, one-way ANOVA, n=4-6).
